# The Theta Paradox – Delayed Reduction of Frontal Midline Theta Following Downregulation Neurofeedback Training

**DOI:** 10.64898/2026.02.03.703090

**Authors:** Thomas Kanatschnig, Lisa Maria Berger, Norbert Schrapf, Markus Tilp, Silvia Erika Kober

## Abstract

Phasic increase of frontal midline theta (Fm theta) has been described as a key indicator of cognitive processing, while relatively lower task-related Fm theta is associated with reduced cognitive strain, reflecting less intensive cognitive processing. In a previous investigation, reduced task-related Fm theta in relation to higher expertise, as well as higher setting anticipation performance in the domain of volleyball was identified. In the present study a single-session sham-controlled neurofeedback training (NFT) intervention was conducted to investigate the feasibility of Fm theta downregulation for the improvement of volleyball setting anticipation. A total of 24 volleyball novices was allocated to “Real” (*n* = 12) and “Sham” (*n* = 12) Fm theta downregulation NFT groups. NFT-related Fm theta, pre-/post-NFT setting anticipation task performance and task-related Fm theta, as well as resting EEG activity were analyzed. Incongruous with our expectations, the Real NFT group showed a tendency toward stronger Fm theta synchronization compared with the Sham group during NFT. Anticipation task performance did not change significantly from before to after NFT in both groups, yet a significant reduction of task-related Fm theta was observed in the Real NFT group following NFT. A post-NFT rebound of Fm theta could be responsible for this result. With our findings we provide further evidence for the existence of an apparent paradox of Fm theta downregulation, in which cognitive control mechanisms, associated with oscillatory Fm theta activity, appear to hinder explicit downregulation of Fm theta through classical neurofeedback learning mechanisms.

## Introduction

Neurofeedback training (NFT) is a prominent technique used by researchers across many different fields – from educational to clinical settings – to investigate neurophysiological substrates of human behavior (Gruzelier, 2014; Ros et al., 2020; Sitaram et al., 2017). NFT can be described as a closed-loop system that combines the measurement of brain activity with real-time sensory (e.g., visual, auditory, tactile) feedback of specific components of neural activation, allowing a person the voluntary manipulation of their own brain activity. Through NFT paradigms, researchers aim to develop new applications for the improvement of various cognitive functions. For that electroencephalography (EEG) has emerged as the de facto standard measurement modality in NFT research, thanks to its high temporal resolution, broad spectrum of neural modulation capabilities, as well as easy and flexible application (Müller-Putz, 2020). Through EEG-based NFT, it is possible to self-regulate oscillatory fluctuations in specific brain frequency bands, such as the theta (4-7 Hz) or alpha (8-12 Hz) bands, which stem from synchronous neuronal activity that can be recorded non-invasively from a person’s scalp (Sitaram et al., 2017). Different frequency bands are associated with different mental processes, but certain mental processes can also generate complementary patterns across multiple frequency bands. Moreover, the spectral power level (i.e., signal intensity) of each frequency band is differently distributed across the cortical surface (Capilla et al., 2022), as well as influenced by psychological (Cohen Kadosh & Staunton, 2019) and anatomical factors (Allison & Neuper, 2010). Also, substantial interindividual differences exist, regarding the ability and strategies used to successfully control one’s own brain activity using NFT (Kober et al., 2013; Witte et al., 2013).

Besides clinical applications, training programs aimed at the improvement of cognitive abilities are one of the most investigated research subjects involving NFT (Gruzelier, 2014). One major area in this regard concerns learning and memory, where it was found that the EEG theta band plays a crucial role in information consolidation and retrieval processes (Gevins et al., 1997; Klimesch, 1999; Lee et al., 2005; Maurer et al., 2015). Klimesch (1999) first described the interconnection between task-/event-related theta and alpha activity, which is characterized by a phasic increase in theta power accompanied by a simultaneous decrease in alpha power during cognitive/memory performance. He hypothesized that phasic increases of theta reflect memory consolidation processes originating from hippocampo-cortical feedback loops (note that from this point onward when referring to theta activity, we generally refer to task-/event-related, i.e., phasic, changes of theta, except when explicitly indicated otherwise). Gevins et al. (1997) demonstrated a positive relationship between increased memory load and frontal midline theta (Fm theta) power – so named due to its primary occurrence over the frontal midline cortical area (Capilla et al., 2022) – further underpinning the involvement of theta in memory consolidation processes. Studies utilizing signal source-localization revealed the anterior cingulate cortex (ACC) as the main source of the Fm theta rhythm (Gevins et al., 1997; Sauseng et al., 2007). Being part of the Papez circuit, the ACC shows interconnections with brain regions involved in learning and memory, such as the amygdala and hippocampus. More recently, Fm theta was identified as a potential key mechanism of cognitive control and attention (Cavanagh & Frank, 2014; Eisma et al., 2021; Kaiser & Schütz-Bosbach, 2021). Given its multifaceted involvement in learning, memory, and cognitive control processes, Fm theta has become a promising target frequency for NFT interventions (Enriquez-Geppert et al., 2014).

In a recent meta-analysis on Fm theta NFT (Pfeiffer et al., 2024) it has been reported that previous studies primarily utilized NFT protocols aiming at Fm theta upregulation, to elicit positive effects on performance measures related to memory (Enriquez-Geppert et al., 2014; Reis et al., 2016; Rozengurt et al., 2017), as well as motoric and executive functions (Rozengurt et al., 2016; Shtoots et al., 2021). The authors conclude that although current research shows promising results on the efficacy of Fm theta NFT for memory enhancement, substantial heterogeneity across studies prevents firm conclusions regarding its true effect size.

Previous studies have also investigated the use of NFT in the domain of sports, where cognitive interventions are an ever-growing research subject (Cheron et al., 2016; Moran, 2009; Seidel-Marzi & Ragert, 2020; Williams et al., 2018). Besides theta and alpha, previous studies commonly utilized the sensorimotor rhythm (12-15 Hz) for NFT, due to its association with motor control and attention, making it a promising target frequency for sports-related interventions. However, as meta-analyses on the subject show, scientifically robust findings regarding the effectiveness of NFT for the improvement of sports-specific cognitive functions are still rare and it is necessary to follow new approaches (Onagawa et al., 2023; Xiang et al., 2018; Yu et al., 2025). In a study by Chen et al. (2022) an Fm theta NFT intervention with the aim of improving golf putting performance was utilized. The authors found that putting performance significantly increased from before to after a single-session of Fm theta NFT in a group of skilled golfers who received function-specific NFT instructions (“*…gradually decrease conscious effort on your action…*”). This improvement was not present in two reference groups receiving different instructions and NFT protocols. More importantly though, the authors conducted Fm theta *downregulation* NFT, which stands in contrast to most related studies utilizing Fm theta upregulation. The study by Chen et al. (2022) was based on a previous study that showed promising preliminary results on a positive effect of Fm theta downregulation on golf putting performance (Kao et al., 2014). Furthermore, the improvement of putting performance in the group that received function-specific instructions occurred alongside a significant reduction of Fm theta from the first to the last block of NFT, while no significant change of Fm theta was observed in the two reference groups.

With the present study, we aimed to expand the evidence on cognition-centered NFT applications in sports by conducting a preliminary investigation into the feasibility of Fm theta downregulation to enhance anticipation performance. This study is based on a study by Kanatschnig et al. (2025), where the neural efficiency hypothesis was tested in the context of volleyball decision-making/anticipation on three groups with differing levels of volleyball expertise, i.e., novices, amateurs and experts. That study described a linear trend between volleyball expertise and a task-related decrease in Fm theta power, which was accompanied by better performance in a video-based *volleyball setting anticipation task* (VSAT). These results were in line with previous findings regarding task-related theta power differences between highly skilled and less skilled athletes (Filho et al., 2021; Li & Smith, 2021). According to a meta-analysis by Filho et al. (2021), higher skilled or expert athletes tend to show lower task-related theta power compared to lesser skilled or novice athletes during sports-specific performance tasks. It is assumed that, due to the positive relationship between Fm theta power and cognitive processes (Gevins et al., 1997; Klimesch, 1999) a lower level of Fm theta could signify a state of lower cognitive strain, or a less “busy” brain, of experts compared to novices during performance. Based on that conclusion we assumed that similar to the positive effects seen for golf putting performance (Chen et al., 2022; Kao et al., 2014), Fm theta downregulation NFT could potentially produce positive effects for volleyball setting anticipation performance as well.

We conducted a single-session sham-controlled NFT intervention for Fm theta downregulation to investigate NFT-induced effects on the setting anticipation ability of volleyball novices, utilizing a pre-/post-test design with an adapted version of the anticipation task for volleyball, previously used by Kanatschnig et al. (2025). Based on the current evidence from investigations on golf putting performance (Chen et al., 2022; Kao et al., 2014) we expected to see positive effects on performance in volleyball setting anticipation, for novices receiving real Fm theta downregulation NFT, but not for novices receiving sham NFT (presentation of prerecorded feedback). Our hypothesis was that novices who received real NFT would show relatively lower task-related Fm theta power during the VSAT after NFT, compared to novices who received sham NFT. Concerning changes of Fm theta power during NFT, we expected the group receiving real NFT to show a linear decline of Fm theta power, while the group receiving sham NFT should show no linear change of power, across six training runs. Therefore, we hypothesized that real Fm theta downregulation NFT would induce stronger Fm theta reduction, i.e., a more negative Fm theta power slope compared to sham NFT.

## Materials and Methods

The procedures and methods for this study were preregistered at the Open Science Framework (OSF; https://osf.io/hzg9j). In Table S1 in the Supplementary Material, we present and discuss deviations from preregistered methods, following the guidelines by Lakens (2024). Furthermore, to comply with the guidelines by Ros et al. (2020) we utilized the “Consensus on the reporting and experimental design of clinical and cognitive-behavioural neurofeedback studies” (CRED-nf) checklist to evaluate our methodological approach for this study. The file “CRED-nf-Checklist.pdf“ provides the complete checklist for the present study, which was created using the official online template and is retrievable from the OSF project page of this study (https://osf.io/bu924/; Kanatschnig et al., 2024).

### Participants

In total 25 participants were recruited for this study. One participant had to be excluded from the analysis due to EEG recording issues during NFT measurements, leaving the data of 24 participants. Among them were 16 women and 8 men, ranging from 19 to 39 years of age (*M* = 25.17 years, *SD* = 4.51). As this study builds on a previous investigation of volleyball anticipation ability (Kanatschnig et al., 2025), we applied the same recruitment criteria used in that earlier study. For the present study only novice participants were recruited, which were defined as individuals with only occasional volleyball experience. Individuals who were actively and regularly playing volleyball at an amateur or professional level in an organized club at the time of data collection were excluded from participation. Participation was still allowed if a person had previously played on a volleyball team or club (as was the case for one participant), or if they had attended a beginner-level volleyball course offered by the local University Sports Institute (USI), which none of the participants had. All participants gave informed written consent before participation. Recruitment and data collection were performed from March 18, 2024, until November 7, 2024. All procedures have been conducted in accordance with the Declaration of Helsinki and were approved by the ethics committee of the University of Graz (GZ. 39/3/63 ex 2023/24).

For our sham-controlled NFT design, participants were allocated to two equally-sized and gender-matched groups, namely the “Real” (age: *M* = 24.00 years, *SD* = 5.24) and the “Sham” (age: *M* = 26.33 years, *SD* = 3.47) group. Each group consisted of twelve participants, we based our sample size rationale on the study by Chen et al. (2022) who recruited twelve participants per NFT group. To examine whether the groups were comparable regarding general sociodemographic variables, as well as volleyball-specific and other sports-related habits (e.g., prior volleyball experience, weekly amount of exercise, …), we calculated comparative statistics for relevant control variables. Furthermore, the Edinburgh Handedness Inventory (Oldfield, 1971) was administered to assess hand dominance. Our analysis did not yield significant results for any control variables, which we interpret as an indication that our NFT groups were sufficiently balanced (see Table S2). Questionnaire presentation was conducted using LimeSurvey (LimeSurvey GmbH).

### Volleyball Setting Anticipation Task

The main objective of our experiment was to investigate task-related changes in Fm theta power during volleyball setting anticipation, in response to an NFT intervention aimed at the downregulation of Fm theta activity. Our domain of interest was volleyball. We used an adapted version of the VSAT used in Kanatschnig et al. (2025). In that earlier publication, the task was referred to as a “tactical decision-making task”; however, it was subsequently renamed to “volleyball setting anticipation task” to provide a conceptually more precise description. The stimulus material of the task consisted of 38 unique video recordings, each depicting a typical 6 vs. 6 women’s indoor volleyball setting scenario and were first used by Schrapf et al. (2022). The videos varied between 3 and 13 s in length and were recorded with a resolution of 720x576 px (5:4 aspect ratio) and frame rate of 25/s during official games of the 2nd Austrian Women’s Volleyball League. In each video, the *server* (player who initiates the rally by serving the ball to the opposing team) of one team prepared for and played the service. After the opposing team received the ball, the *setter* (player who delivers the final pass, i.e., the setting/set, for the ensuing attack) got into position to set up the attack. All videos were edited to stop 0.08 seconds (two frames) before the setter touches the ball, so that the setter’s pass destination was occluded. Following each video, participants were asked to make a prediction on the outcome of the shown setting sequence, namely to which court position the setter will pass the ball (“*Where will the SETTER pass the ball to?*”).

The presentation of each video stimulus followed the same structure. Before the onset of each video, participants had to fixate their gaze on a fixation disk in the center of the screen for 2 s in the “Baseline” phase. The time interval from the start of the video until the server played the ball marks the “Preparation” phase, which lasted between 0 to 10 s (variation in length was due to differing preparation times of the server in each video). The Preparation phase was followed by the “Rally” phase, which was the main phase of interest in our analysis. It was set to a fixed length of 3 s and included the course of the rally from the service until the end of the video (due to the consistent structure of each rally sequence, 3 s was consistently found to be the time frame between service and setting). The Rally phase was followed by the “Freeze” phase, in which the last video frame remained visible on screen for 0.5 s, after which a response screen appeared, prompting participants to make their prediction regarding the setter’s pass destination in the “Response” phase. For a visualization of the procedure during the VSAT see Fig. 1A^1^. The task was implemented using PsychoPy (ver. 2022.1.2; Peirce et al., 2019).

**Fig. 1.**
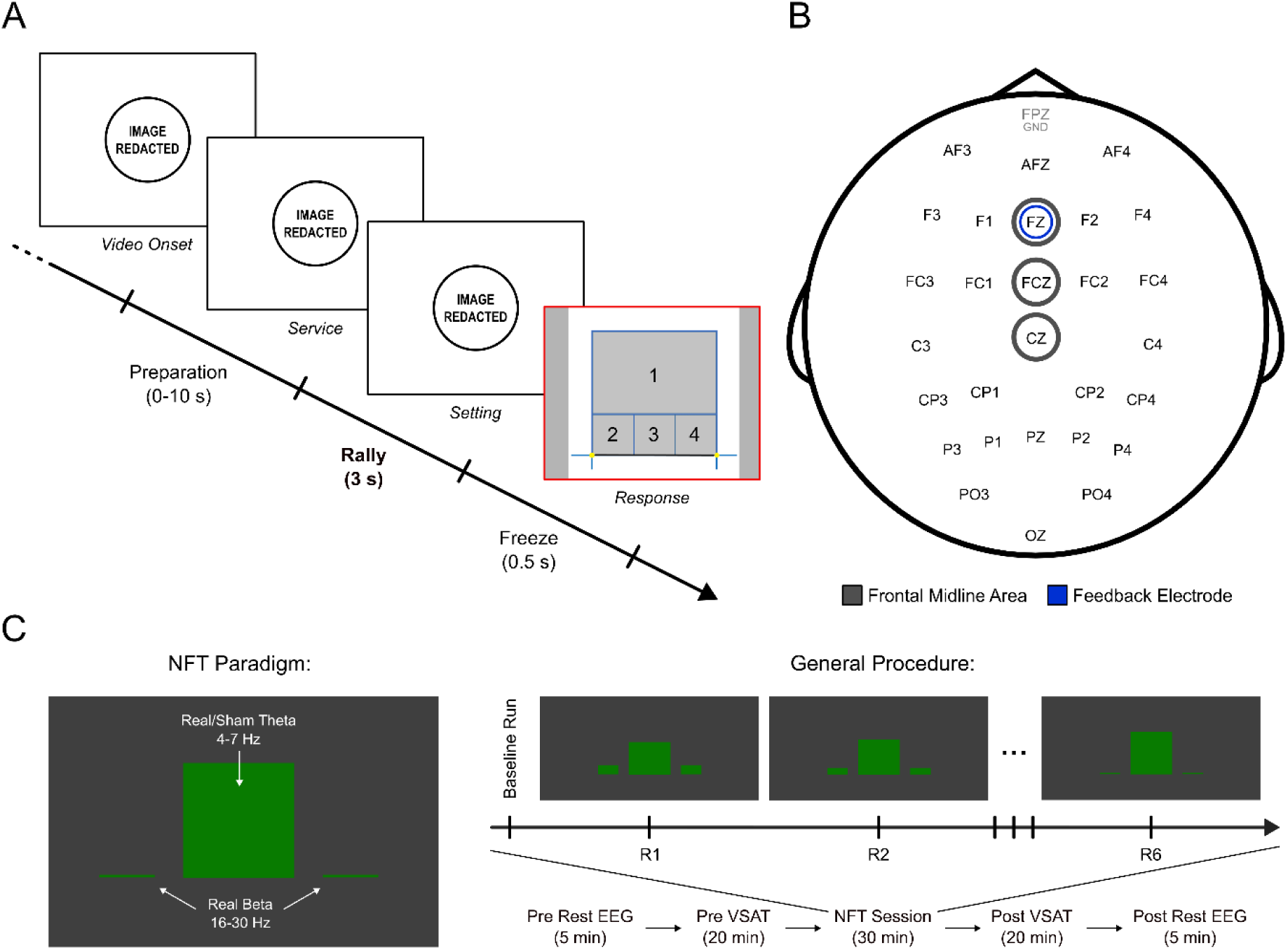
Visualization of experimental procedure and EEG montage. (A) volleyball setting anticipation task procedure (the main task phase “Rally” is marked in bold font), (B) EEG electrode locations according to EEG 10-10 system (“GND”: ground electrode), and (C) neurofeedback training paradigm and general procedure (“NFT”: neurofeedback training; “VSAT”: volleyball setting anticipation task). Image material partially reprinted from Kanatschnig et al. (2025) under a CC BY license, original copyright 2025.

For the investigation of NFT-related effects on volleyball setting anticipation ability, we incorporated a pre-/post-test design, in which we presented two comparable subsets of video stimuli to participants, before and after NFT, respectively. The two task versions “A” and “B” each included 19 video stimuli, which were selected based on the results of the previous study (Kanatschnig et al., 2025), i.e., both task versions included a subset of stimuli that were comparable in terms of difficulty (prediction accuracies in the novice sub-group from the previous study) and visual properties (all videos depicted game sequences from the same camera perspective).

### Neurofeedback Training

Our single-session sham-controlled NFT intervention comprised one “Baseline” run followed by six main NFT runs, in which a basic visual bar paradigm was presented. The Real group received simultaneous feedback of their real theta (4-7 Hz) as well as beta (16-30 Hz) activity, while the Sham group received only real beta feedback and sham theta feedback, which was the prerecorded visual feedback from a gender-matched participant of the Real group. The FZ electrode functioned as the feedback electrode for both theta as well as beta activity. See Fig. 1B for a visualization of the EEG montage including the frontal midline region of interest (ROI). The Baseline and NFT runs each lasted approx. 3 min (measuring start and stop was performed manually). The NFT paradigm was programmed and presented using Unity (ver. 2020.3.30f1; Unity Technologies), data streaming between Unity and EEG was done via Lab

Streaming Layer (LSL) and the *LSL4Unity* plugin (https://github.com/jelenaLis/LSL4Unity). Participants saw three green bars centered in the middle of the screen in front of a grey background; one larger bar in the middle and two smaller bars on the sides. During the Baseline run, the three bars moved up and down continuously without any connection to participants’ brain activity, to measure baseline theta and beta activity for individual thresholding. Participants were asked to follow the bar movement visually, without any specific mental task. Baseline theta (median) and beta (median + 1 mean absolute deviation) activity were set as thresholds for all six NFT runs. For the NFT runs, participants were instructed to “*… find your own strategy to keep the bar in the middle as high as possible and at the same time keep the bars on the sides as low as possible.*” Participants were also told that it is possible to change strategies at any time during training and to try to avoid muscle tensions and reduce eye blinks. During NFT runs, the middle bar represented theta activity, while the outer bars represented beta activity. As previously mentioned, the aim of the NFT session was to downregulate theta activity. The direction of the theta bar was inverted so that it increased when theta activity reduced and vice versa, as we deemed it to be more natural for participants to try increasing the middle bar instead of reducing it. Besides the main aim of increasing the theta bar, the two beta bars functioned as a visual artifact control. The bars increased when beta activity increased, making the participants aware of artifact interference. For a visualization of the NFT procedure see Fig. 1C.

### EEG Recording and Preprocessing

The EEG montage used in this study consisted of 28 scalp electrodes that were arranged following a placement layout, aimed specifically at the coverage of frontal as well as parieto-occipital regions (see Fig. 1B). Reference electrodes were placed on the right and left mastoids. The ground electrode was placed on the FPZ location. Three EOG electrodes were used, which were placed according to Schlögl et al. (2007). The actiCAP slim active electrode system was used in conjunction with BrainAmp amplifiers (BrainProducts GmbH). During VSAT measurements EEG was recorded using BrainVision Recorder (ver. 1.26.0101; BrainProducts GmbH), during NFT measurements EEG was recorded and processed using OpenViBE (ver. 3.5.0; Renard et al., 2010). Online feature extraction during NFT was based on the procedures applied by Berger et al. (2025). Time-based epoching was performed using 1 s epochs with 0.5 s overlaps. The data was squared to obtain power values, which were then averaged. Following the application of a moving average across epochs, the data was natural log-transformed and streamed to Unity via LSL for implementation in the feedback loop.

The recorded EEG data was preprocessed by means of the following processing steps within BrainVision Analyzer (ver. 2.3.0.8300; BrainProducts GmbH). A linked-mastoid reference was formed from the two reference channels and applied to all channels. To eliminate low-frequency drift and power line interference a 0.5 Hz low cutoff as well as a 50 Hz notch filter was applied. Manual artifact correction was performed, in which larger muscle artifacts were cut. After artifact correction a 70 Hz high cutoff filter was applied to eliminate high-frequency noise. Lastly, eye movement artifacts were identified and corrected using independent component analysis, which was followed by semi-automatic artifact exclusion based on the following EEG criteria: (i) > 50 μV voltage difference between two data points, (ii) > 200 μV voltage difference within a 200 ms interval, and (iii) absolute voltage values ±120 μV. Signal that fell under one of these criteria was marked 500 ms before and after the identified datapoints. Remaining artifacts and EEG channels with high noise levels were excluded manually. Following artifact correction, extraction of theta (4-7 Hz) and beta (16-30 Hz) frequency band power (µV²) was performed using the *Complex Demodulation* transformation function within BrainVision Analyzer. In the case of VSAT, segmentation and averaging of EEG power data was performed for the Baseline phase as well as the main task phase of interest, the Rally phase. In the case of NFT, EEG power data was cut into 2 s segments and averaged across each run. The same segmentation and averaging procedure was performed for resting EEG data. The averaged EEG power data was then further processed using R (ver. 4.4.3; R Core Team, 2021), RStudio (ver. 2025.05.0+496; RStudio Team, 2020) and the *tidyverse* package (ver. 2.0.0; Wickham et al., 2019). EEG theta and beta power data was then transformed into event-related de-/synchronization (ERD/S) data using the following formula:

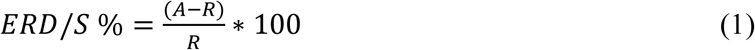

In this formula (Pfurtscheller & Lopes da Silva, 1999) “A” stands for an *active* period (i.e., VSAT phase, NFT run), while “R” stands for the respective *resting* period (i.e., Baseline phase/run). Positive theta and beta values resulting from this transformation signify an event-related synchronization (ERS), while negative values signify an event-related desynchronization (ERD). Due to the standardization of signal levels across participants, the ERD/S technique provides improved comparability for NFT-related and task-related EEG signal changes.

### General Procedure

Data acquisition for this study was performed in an electrically shielded and sound attenuated EEG laboratory at the University of Graz. After providing written consent, participants filled out a questionnaire including sociodemographic and control variables. Following that, the EEG montage was applied and prepared for measurement. Next, resting EEG measurements were performed with open as well as closed eyes for 2.5 min each, followed by the instruction for the VSAT. Participants then performed a short practice sequence to familiarize themselves with the task, which included different stimuli from the ones in the main task measurements. Then, the first measurement of the VSAT was performed. This was followed immediately by the NFT intervention, which was followed by the second measurement of the VSAT. The presentation order of VSAT versions “A” and “B” was counterbalanced across participants. After that a second resting EEG measurement was performed, analogous to the first one. Finally, participants completed a short questionnaire including visual analog scale (VAS) variables that assessed subjective ratings of factors such as motivation, concentration, and perceived performance during the experiment. Comparative statistics for VAS variables did not reveal any significant group differences (see Table S3). Participants also answered open questions regarding the choice of strategies during NFT; a qualitative comparison of participants’ NFT strategies did not reveal any specific group differences (see Table S4). A diagram of the general experimental procedure is provided in Fig. 1C (not depicted are pre- and post-experiment questionnaire assessments).

### Dependent Variables

The main dependent variables (DV) of our analyses consisted of behavioral performance measures during the VSAT, as well as neurophysiological measures during the VSAT and during NFT. Behavioral performance during the VSAT was defined as the response accuracy, i.e., the percentage of correct responses given for the items of the task, to which we refer to as “Task Accuracy”. We additionally investigated “Task Response Time”, i.e., the average response time participants took to give their response. Regarding neurophysiological DVs, we investigated Fm theta ERD/S during the completion of the VSAT, i.e., “Task Fm Theta ERD/S”, as well as during NFT, i.e., “NFT Fm Theta ERD/S”. In the case of Task Fm Theta ERD/S the analyzed EEG activity was that of the Rally phase of the task, for which ERD/S was derived from the above described transformation (see section EEG Recording and Preprocessing), in which the Rally phase activity was corrected by the activity during the Baseline phase. For NFT Fm Theta ERD/S the baseline-corrected activity during each main NFT run was analyzed. On an exploratory basis, we also investigated pre- and post-experiment resting Fm theta activity, i.e., “Rest Fm Theta”, for which we present the analysis of eyes-open resting EEG, due to its higher comparability with our active task and NFT paradigms. Analogous to Kanatschnig et al. (2025) we used theta ERD/S [%] from frontal midline electrodes FZ, FCZ and CZ for the analysis of Task Fm Theta ERD/S and NFT Fm Theta ERD/S, while for the analysis of Rest Fm Theta, we used natural log-transformed theta power [log(µV²)] from the same frontal midline electrodes (see Fig. 1B).

### Statistical Analysis

For the investigation of NFT related effects on behavioral Task Accuracy and Task Response Time (natural log-transformed average response times were used for statistical analysis), we calculated linear mixed effect models (LMM), including the fixed factors *Group* (Real vs. Sham) and *Measurement* (Pre vs. Post), as well as a random intercept term (1 | *Code*) to account for general baseline differences between participants (represented by their participation Code). For the investigation of Task Fm Theta ERD/S the LMM structure was expanded by the fixed factor *Electrode* (FZ vs. FCZ vs. CZ). The analysis of a group difference for NFT outcome was conducted by means of an independent samples *t*-test including the factor *Group* as independent variable and the slope of NFT Fm Theta ERD/S across the six NFT runs as DV (degrees of freedom were adjusted by means of the Welch method; Welch, 1947). To obtain NFT slopes, simple linear regressions were calculated for each participant’s NFT performance, including the predictor variable *Run* (R1-R6) and the outcome variable NFT Fm Theta ERD/S. For control purposes we performed the same calculations also for the beta band. For the investigation of pre- and post-experiment Rest Fm Theta, the same LMM structure as with Task Fm Theta ERD/S was used. The following formula depicts our LMM structure:

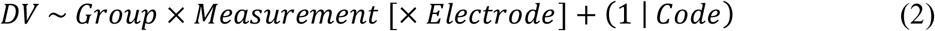

LMMs were calculated by means of the *lme4* (ver. 1.1-36; Bates et al., 2015) and *lmerTest* (ver. 3.1-3; Kuznetsova et al., 2017) packages for R. Global fixed effects were tested with a Type III ANOVA for each model. To identify group differences we calculated pairwise comparisons using the *emmeans* (ver. 1.11.0, Lenth, 2023) package. The *rstatix* (ver. 0.7.2.; Kassambara, 2023b), *car* (ver. 3.1-3; Fox & Weisberg, 2019) and *e1071* (ver. 1.7-16; Meyer et al., 2024) packages were additionally used for statistical calculations. The *eegUtils* (ver. 0.8.0; Craddock, 2022), *ggpubr* (ver. 0.6.0; Kassambara, 2023a), *patchwork* (ver. 1.3.0; Pedersen, 2025) and *sjPlot* (ver. 2.8.17; Lüdecke, 2024) packages were used for visualization purposes. Degrees of freedom for LMMs were calculated using Satterthwaite’s method (Satterthwaite, 1946). Effect sizes provided for pairwise comparisons were computed using the *eff_size* function of *emmeans* and can be interpreted as an estimation of Cohen’s *d*. The significance level for all analyses was set to α = 0.05 (two-tailed). When calculating multiple pairwise comparisons, the significance level was automatically adjusted using the Holm method (Holm, 1979). All main, exploratory, as well as complementary analyses, including model diagnostics, can be examined using the respective data files and R code retrievable from this study’s OSF project page (https://osf.io/bu924/; Kanatschnig et al., 2024).

## Results

Our hypothesis stated that an overall negative linear trend of NFT Fm Theta ERD/S over the course of the six NFT runs was expected for the Real feedback group, while no specific trend was predicted for the Sham group. Conversely, the subject-level regression analyses revealed that the majority of Sham group participants (9/12) showed a negative NFT slope, while the opposite was the case in the Real group, where more participants (7/12) showed a positive, instead of a negative NFT slope. Yet, regarding average NFT slopes, the two groups did not differ substantially (Real: *M* = 0.80, *SD* = 5.78; Sham: *M* = 0.19, *SD* = 2.78). Consequently, a *t*-test for a group difference in NFT slopes showed a non-significant result (*t*(15.8) = 0.33, *p* = .747, *d* = 0.13). Results for the analysis of NFT performance are presented in Fig. 2 for each group, respectively. Fig. 2A shows NFT performance of all participants separated by NFT slope direction (+/-), while Fig. 2B shows boxplots for each run of the NFT intervention. The analysis of beta activity did not reveal significant group differences, implying that there were no substantial differences in EEG artifact production between groups (beta analysis results can be inspected through the provided code “eeg_nft_theta+beta_erds_analysis.R” in the online materials, alongside all other main statistical analysis files; Kanatschnig et al., 2024).

**Fig. 2.**
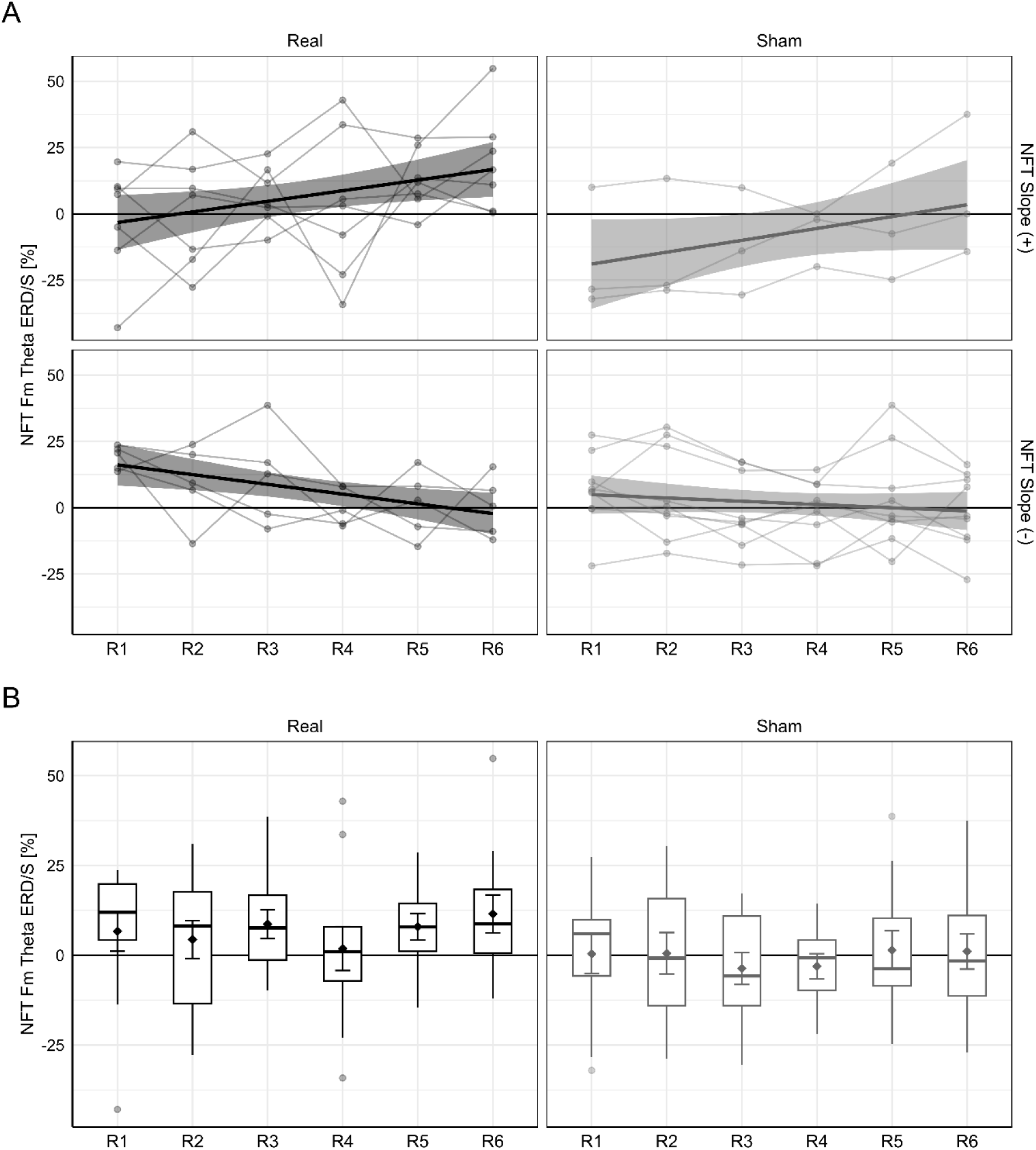
Results for the analysis of NFT Fm Theta ERD/S. (A) NFT performances from individual participants of each NFT group (Real, Sham) over the course of the six runs (R1-R6) are represented by line-connected dots and seperated by NFT slope direction. Regression lines and confidence ribbons represent linear regression calculations for each figure panel. (B) Boxplots of group NFT performance for each run (R1-R6). Means and standard errors are represented within boxplots by diamond symbols and error bars.

Following our hypotheses regarding NFT-induced changes in behavioral task performance, as well as task-related Fm theta power, our expectation was to find significant pre/post differences in the respective LMM analyses. The full ANOVA results from all LMM analyses are provided in Table 1.

**Table 1.** ANOVA results for LMM analyses.

| **DV:** | **Fixed Effect:** | ***SumSq*** | ***MeanSq*** | ***df*** | ***F*** | ***p*** | ***Sig.*** |
| --- | --- | --- | --- | --- | --- | --- | --- |
| Task Accuracy | Group | 8.22 | 8.22 | 1, 22 | 0.07 | .789 |  |
|  | Measurement | 97.53 | 97.53 | 1, 22 | 0.87 | .362 |  |
|  | Group × Measurement | 129.85 | 129.85 | 1, 22 | 1.15 | .294 |  |
| Task Response Time | Group | 0.04 | 0.04 | 1, 22 | 1.90 | .182 |  |
|  | Measurement | 0.08 | 0.08 | 1, 22 | 3.29 | .083 |  |
|  | Group × Measurement | 4.30e-03 | 4.30e-03 | 1, 22 | 0.18 | .673 |  |
| Task Fm Theta ERD/S | Group | 74.44 | 74.44 | 1, 22 | 0.33 | .570 |  |
|  | Measurement | 2329.34 | 2329.34 | 1, 110 | 10.41 | .002 | ** |
|  | Electrode | 53.73 | 26.86 | 2, 110 | 0.12 | .887 |  |
|  | Group × Measurement | 1531.56 | 1531.56 | 1, 110 | 6.84 | .010 | * |
|  | Group × Electrode | 90.17 | 45.08 | 2, 110 | 0.20 | .818 |  |
|  | Measurement × Electrode | 98.87 | 49.44 | 2, 110 | 0.22 | .802 |  |
|  | Group × Measurement × Electrode | 14.62 | 7.31 | 2, 110 | 0.03 | .968 |  |
| Rest Fm Theta | Group | 3.52e-03 | 3.52e-03 | 1, 22 | 0.10 | .761 |  |
|  | Measurement | 0.01 | 0.01 | 1, 110 | 0.32 | .574 |  |
|  | Electrode | 0.61 | 0.30 | 2, 110 | 8.23 | < .001 | *** |
|  | Group × Measurement | 0.17 | 0.17 | 1, 110 | 4.47 | .037 | * |
|  | Group × Electrode | 0.12 | 0.06 | 2, 110 | 1.60 | .207 |  |
|  | Measurement × Electrode | 9.95e-03 | 4.98e-03 | 2, 110 | 0.13 | .874 |  |
|  | Group × Measurement × Electrode | 8.83e-06 | 4.42e-06 | 2, 110 | 0.00 | .999 |  |

*Note.* Type III ANOVA results with degrees of freedom calculations using Satterthwaite's method. Significant fixed effects are indicated with asterisks (\*: $p < .05$ ; \*\*: $p < .01$ ; \*\*\*: $p < .001$ ).

On the behavioral level, LMMs for Task Accuracy and Task Response Time did not show any significant main or interaction effects. Nonetheless, as the modulation of volleyball setting anticipation performance through NFT was one of the main variables of interest of our investigation, a visualization of the performance results for Task Accuracy is provided in Fig. 3A. Descriptive statistics for pre-/post-NFT comparisons are provided in Table 2 (for subject-level statistics see Table S4).

**Fig. 3.**
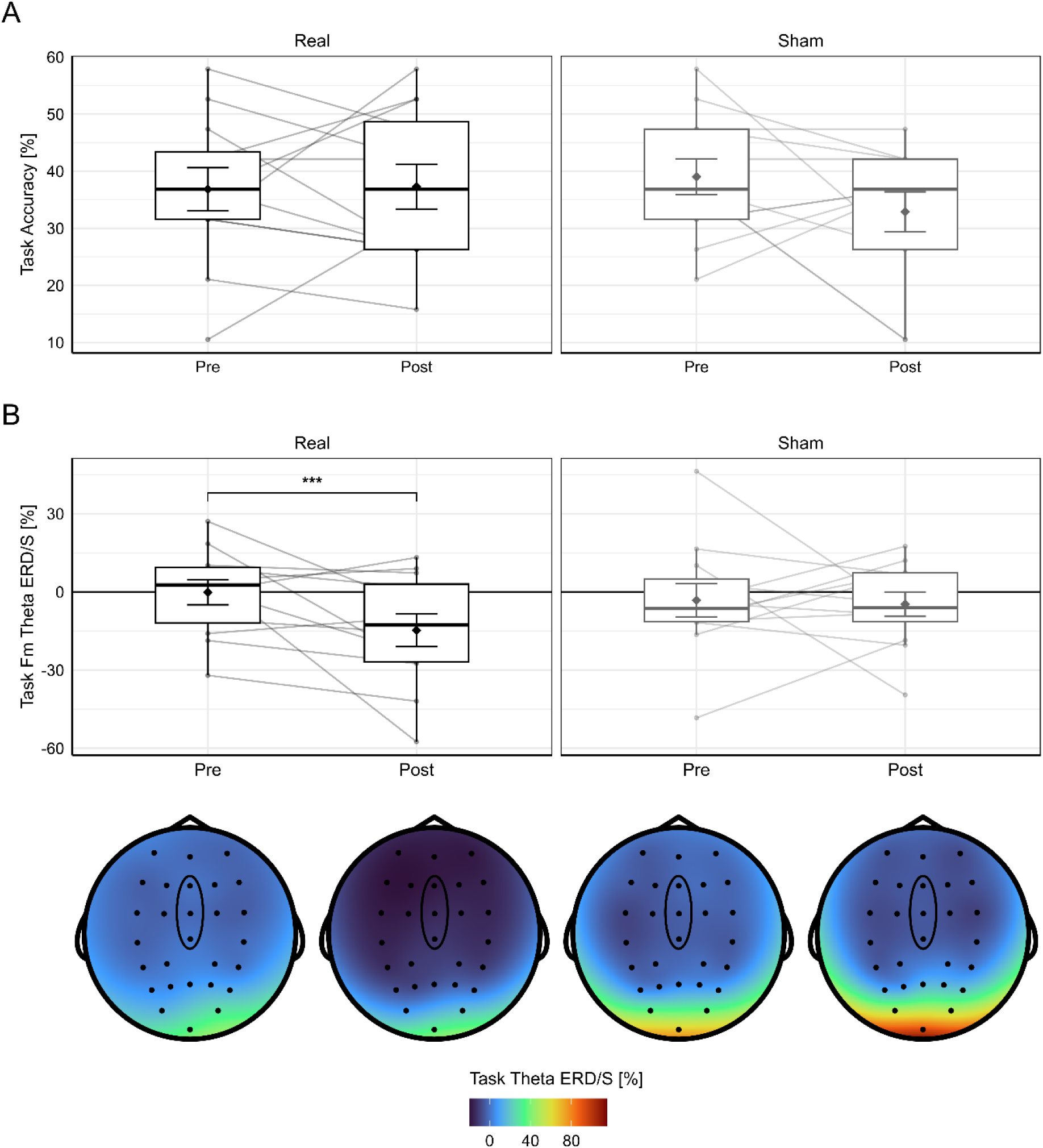
Results for the analysis of (A) Task Accuracy, and (B) Task Fm Theta ERD/S. Pre/post differences of individual participants of each NFT group (Real, Sham) are represented by line-connected dots. Means and standard errors are represented within boxplots by diamond symbols and error bars. Significant pre/post differences, as determined through LMM analysis, are indicated with asterisks (***: *p* < .001). Corresponding topographical plots are presented below each Task Fm Theta ERD/S boxplot (note that negative dark blue values indicate a task-related power decrease, i.e., ERD).

**Table 2.** Descriptive statistics for pre-/post-comparisons.

| **DV:** | **Real (n = 12):** |  | **Sham (n = 12):** |  |
| --- | --- | --- | --- | --- |
|  | *Pre* | *Post* | *Pre* | *Post* |
| Task Accuracy [%] | 36.84 (±13.09) | 37.28 (±13.55) | 39.04 (±10.87) | 32.90 (±12.11) |
| Task Response Time [s] | 2.68 (±1.09) | 2.42 (±0.97) | 2.14 (±0.61) | 2.05 (±0.72) |
| Task Fm Theta ERD/S [%] | -0.17 (±16.64) | -14.74 (±21.67) | -3.20 (±22.34) | -4.72 (±16.19) |
| Rest Fm Theta [log(μV<sup>2</sup>)] | 3.17 (±0.45) | 3.12 (±0.58) | 3.05 (±0.31) | 3.14 (±0.37) |

*Note.* Means and standard deviations for pre- and post-NFT measurements are presented for each group within the respective group columns.

On the neurophysiological level, the LMM for Task Fm Theta ERD/S revealed a significant interaction effect *Group* × *Measurement*, as well as a significant main effect *Measurement*. No other main or interaction effects for Task Fm Theta ERD/S reached significance. Pairwise comparisons for *Group* × *Measurement* revealed that the Real group showed significant stronger Fm theta ERD (task-related decrease in Fm theta power) during the VSAT after NFT compared to before NFT (*t-ratio*(110) = 4.13, *p* < .001, *d* = 0.72), while there was no significant pre/post difference in Task Fm Theta ERD/S in the Sham group (*t-ratio*(110) = 0.43, *p* = .667, *d* = 0.08). A visualization of Task Fm Theta ERD/S is provided in Fig. 3B, which includes a full topographical representation of task-related theta ERD/S, showing a broad frontocentral decline in theta during the post-measurement of the VSAT in the Real group. Examining this pre/post difference in the Real group at the single-electrode level revealed that the effect is present for all electrodes in the frontal midline ROI (for a visualization see Fig. S1).

The LMM results from our exploratory investigation of pre- and post-NFT Rest Fm Theta activity showed a significant interaction effect *Group* × *Measurement*, as well as a significant main effect *Electrode*. No other main or interaction effects for Rest Fm Theta reached significance. Note that for resting theta activity natural log-transformed theta power [log(µV²)] was analyzed. Despite a significant effect, pairwise comparisons for *Group* × *Measurement* showed no significant pre/post difference in the Real group (*t-ratio*(110) = 1.10, *p* = .275, *d* = 0.11), and a difference which was only on the verge of significance in the Sham group (*t-ratio*(110) = -1.90, *p* = .061, *d* = -0.19). On a topographical level, pairwise comparisons for the factor *Electrode* revealed significantly stronger theta activity at FCZ (*M* = 3.20, *SD* = 0.43) compared to CZ (*M* = 3.04, *SD* = 0.34; *t-ratio*(110) = 4.05, *p* < .001, *d* = 0.35), but not compared to FZ (*M* = 3.13, *SD* = 0.49; *t-ratio*(110) = -1.80, *p* = .075, *d* = -0.16). The difference between FZ and CZ was marginally non-significant (*t-ratio*(110) = 2.25, *p* = .053, *d* = 0.19). A visualization of Rest Fm Theta is provided in Fig. 4 including full topographical representation, showing similar distribution patterns of pre- and post-NFT resting theta activity across groups, with a frontocentral peak around the FZ and FCZ electrodes.

**Fig. 4.**
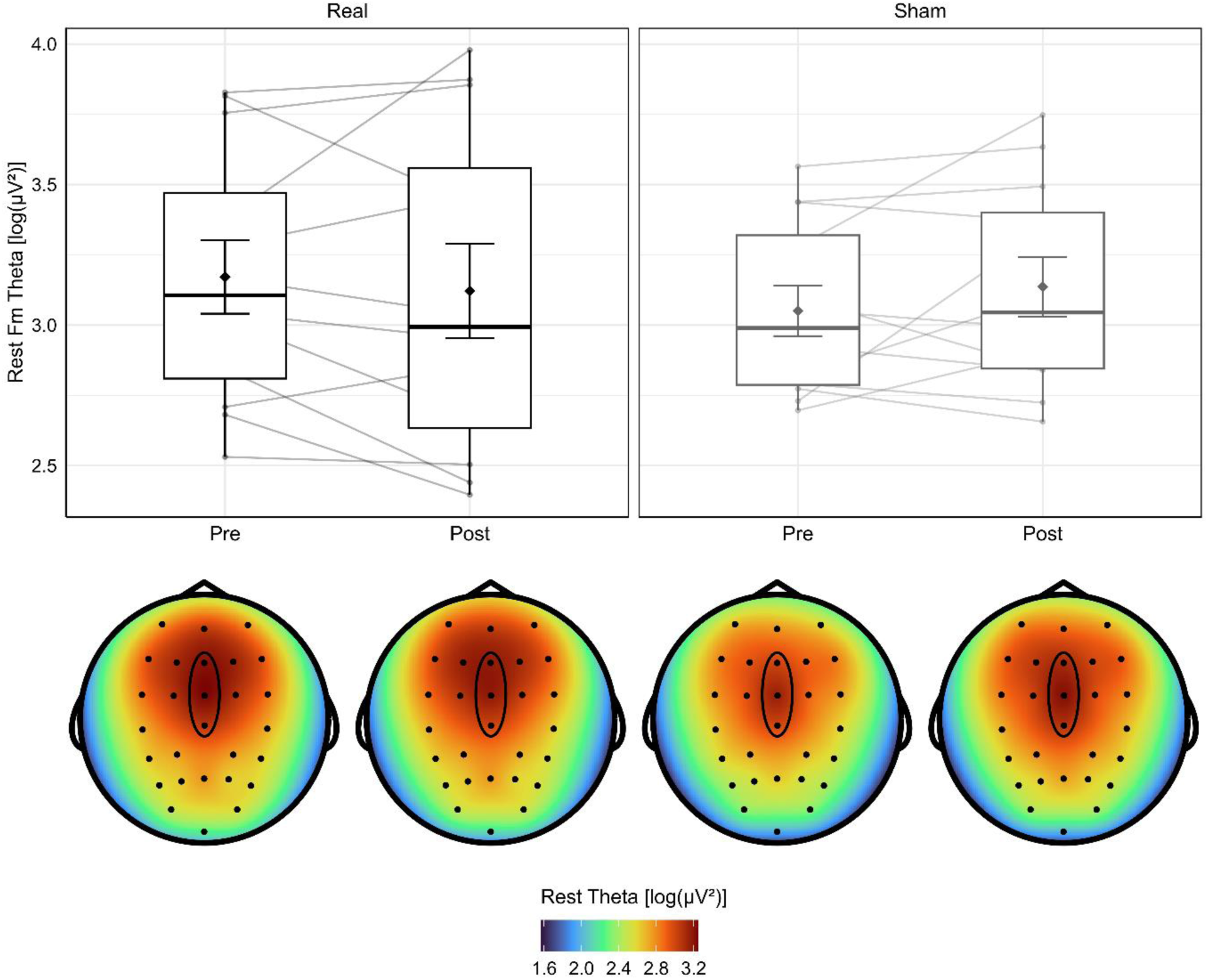
Results for the analysis of Rest Fm Theta. Pre/post differences of individual participants of each NFT group (Real, Sham) are represented by line-connected dots. Means and standard errors are represented within boxplots by diamond symbols and error bars. Natural log-transformed resting theta power was analyzed. Corresponding topographical plots are presented below each Rest Fm Theta boxplot.

## Discussion

In the present study we investigated the effects of Fm theta downregulation NFT on sports-related anticipation performance as well as theta activity. Our sample consisted of volleyball novices who performed a single-session sham-controlled NFT intervention, with the aim of increasing volleyball setting anticipation performance during a video-based task, by reducing Fm theta power (Chen et al., 2022; Kao et al., 2014). While our results did not show changes in behavioral task performance during setting anticipation from pre- to post-NFT measurements, we found a significant decrease in relative Fm theta power from before to after NFT during the task, in the group that received real NFT. While our results revealed the expected reduction of task-related Fm theta power following real NFT, the analysis of NFT-related changes in Fm theta power showed a contradictory pattern, in which the participants of the Real NFT group showed tendentially increased (stronger Fm theta synchronization) instead of reduced Fm theta power during downregulation NFT. Our findings provide notable new evidence on neurophysiological dynamics of Fm theta in the field of Fm theta downregulation NFT.

The results of our analyses present an apparent paradox: task-related Fm theta power is reduced following theta downregulation NFT in the intervention group receiving real NFT, while the majority of participants in that group showed NFT performance slopes in the opposite, *positive*, direction. In contrast to prior research on Fm theta NFT (Pfeiffer et al., 2024) our results stand in conflict with the predominantly reported coherent pattern between the direction of the conducted NFT protocol and Fm theta modulation outcome (i.e., increase after upregulation). In their meta-analysis Pfeiffer et al. (2024) summarized that the majority of investigated studies which utilized upregulation NFT protocols showed successful within-session Fm theta increase (Enriquez-Geppert et al., 2014; Eschmann & Mecklinger, 2022; Rozengurt et al., 2016, 2017; Shtoots et al., 2021). Due to the small number of studies, a clear synthesis regarding the success rate of Fm theta downregulation is not feasible at this moment; however, studies that are currently available have shown peculiar results. Interestingly, studies that utilized theta downregulation NFT as a control group protocol showed overall unsuccessful downregulation performance as well (Rozengurt et al., 2017; Shtoots et al., 2021). Meanwhile, Chen et al. (2022) reported successful within-session reduction of Fm theta in an NFT group receiving a function-specific instruction (“*…gradually decrease conscious effort on your action…*”), but unsuccessful reduction both in an NFT group receiving a traditional instruction (“*…develop your own strategies to control the brain wave…*”) and a control group receiving sham NFT. Kao et al. (2014) conducted a downregulation NFT intervention as well, yet their sample was limited to only three participants and no within-session Fm theta changes were reported.

The question of why our NFT intervention did not induce the intended reduction of Fm theta is difficult to answer. Pfeiffer et al. (2024) concluded that learning theta downregulation appears to be more difficult than learning theta upregulation and therefore a single-session NFT intervention might not be sufficient to induce the intended effect. More importantly though, they identified an unrecognized yet significant dilemma regarding Fm theta downregulation NFT. Fm theta has been shown to play an instrumental role in cognitive control processes (Cavanagh & Frank, 2014; Eisma et al., 2021). This includes aspects such as self-regulation (Ninaus et al., 2013) and reward processing (Amiez et al., 2012), which are key mechanisms of neurofeedback learning (Sitaram et al., 2017). As a consequence, the experimental setting of NFT itself might prohibit the voluntary downregulation of Fm theta. As the focused allocation of mental resources on the presented feedback modality during NFT should produce an increase of Fm theta, this would result in a conflict with the intended goal of downregulating Fm theta. This could explain why, in the study by Chen et al. (2022), only the group receiving the function-specific instruction was able to reduce Fm Theta during NFT, as the authors provided a specific strategy to downregulate conscious effort. This raises the assumption that the reduction of Fm theta in this group potentially occurred due the function-specific instruction – aimed at a voluntary top-down directed reduction of mental effort – rather than due to theta downregulation NFT. Moreover, the group receiving a traditional instruction to find their own strategy for self-regulating Fm theta activity, analogous to our findings, showed no significant NFT-related within-session changes in Fm theta.

If the interpretation that the apparent conflict between feedback processing and Fm theta modulation (Pfeiffer et al., 2024) is further supported by substantive evidence, it would have consequential implications regarding the feasibility of Fm theta downregulation NFT itself. It would suggest that traditional neurofeedback learning paradigms, which are based on operant conditioning (Kamiya, 2011) and reinforcement mechanisms (Siniatchkin et al., 2000), are not suited for Fm theta downregulation. The results of the present study, as well as previous studies involving Fm theta downregulation (Rozengurt et al., 2017; Shtoots et al., 2021) seem to align with this hypothesis; nevertheless, more research is necessary for further validation. There is also the possibility that Fm theta downregulation performance might be improved through applying covert NFT (Muñoz-Moldes & Cleeremans, 2020; Ramot et al., 2016). In an fMRI-based NFT experiment Ramot et al. (2016) demonstrated the possibility of administrating unconscious neurofeedback, meaning that the participants receiving NFT were unaware of the fact that their performance on a given task was being influenced by their own brain activation. In that study the authors concluded that brain networks can be modified through NFT even in complete unawareness of the NFT feedback-loop. We assume that such an implicit covert NFT approach could benefit Fm theta downregulation performance as well, as active attentional control mechanisms of classical NFT learning paradigms are intentionally circumvented in such scenarios. However, it must be considered that the setup for this type of NFT is much more elaborate, as participants must be kept unaware of the fact that (i) they are performing NFT, and (ii) that there is a dependency between the use of specific mental strategies and the performance outcome (Muñoz-Moldes & Cleeremans, 2020).

As our results showed unsuccessful Fm theta downregulation during NFT, yet a significant reduction of task-related Fm theta power followed by real (but not sham) NFT, the question remains how this reduction of Fm theta in the Real group emerged following the NFT intervention. A potential explanation for this phenomenon is the occurrence of a temporally coupled rebound effect in Fm theta, which has previously been described for post-movement beta band activity (Neuper & Pfurtscheller, 2001) as well as following alpha band downregulation NFT (Kluetsch et al., 2014). Ros et al. (2014) explained this phenomenon through the involvement of homeostatic plasticity, which suggests that an initial focal increase in neural activation is followed by a compensatory decrease to maintain organismic homeostasis of neural activation levels. Furthermore, we assume that by being provided with a direct interface for controlling Fm theta through NFT, the participants of the Real group built up a more direct attentional connection with their own theta activity, thereby increasing phasic oscillatory theta activity in general, due to the involvement of Fm theta in attention and cognitive control processes. This is also reflected by the tendentially heightened average ERS levels of the Real group during NFT (see Fig. 2B). As a consequence, this more pronounced Fm theta ERS (stronger increase of Fm theta power) potentially led to a rebound of Fm theta as a form of homeostatic compensation after completion of the NFT intervention, resulting in a stronger Fm theta ERD (stronger decrease of Fm theta power) during the subsequent VSAT.

It is unclear what specific oscillatory dynamics Fm theta underlies in relation to those of other EEG frequencies, yet the rebound of Fm theta following our NFT intervention shows apparent similarities to that of previous reports of the rebound effect (Kluetsch et al., 2014; Neuper & Pfurtscheller, 2001; Ros et al., 2014). Furthermore, concerning the timescale of the effect, our findings suggest that a significant difference in EEG band power levels may be maintained up to several minutes after NFT offset, as the post-NFT measurement of the VSAT lasted approx. 20 minutes. Moreover, the rebound occurred during active task performance, which implies that the effect prolongs not only during passive resting measurements, as found following alpha NFT (Kluetsch et al., 2014), but also during phases of active cognitive engagement. However, we did not identify the same level of Fm theta reduction in the subsequent post-NFT resting measurement, indicating the temporal limitation of the rebound effect. Keeping these constraints in mind, our results provide a new perspective on the use of a transient NFT-induced rebound effect for neural activity modulation.

Finally, although the Real group exhibited a significant reduction in Fm theta power during the VSAT following NFT, this change did not appear to affect setting anticipation performance. The question remains why our NFT intervention did not yield behavioral performance changes comparable to those reported in previous studies on golf putting performance (Chen et al., 2022; Kao et al., 2014), which demonstrated behavioral performance improvements alongside Fm theta modulation. One explanation may be that, whereas those studies focused on skilled golf players, our sample consisted of novices with only limited prior experience in volleyball. It must be considered that higher domain-specific expertise of skilled volleyball players could potentially have affected behavioral task performance as well as NFT performance differently. It is also possible that a primarily passive anticipation task such as the VSAT is not as sensitive to Fm theta modulation as an active task such as golf putting. The modulation of Fm theta activity towards a state of lower cognitive strain (Gevins et al., 1997; Klimesch, 1999) through downregulation NFT could potentially have a more pronounced impact on volleyball setting anticipation performance when investigated in a sample of advanced volleyball players as well as under more ecologically valid circumstances.

## Limitations

Limitations of our study include the circumstance that we did not incorporate a double-blinded design, which has the potential to reduce experimenter bias during NFT interventions (Ros et al., 2020). However, since our instruction and training procedure during NFT were highly standardized, we do not assume that this significantly impacted our results. Another caveat to our findings is the fact that, given the preliminary nature of this study, we only conducted a single-session NFT intervention, therefore providing only a small snapshot of the underlying mechanisms of Fm theta downregulation NFT (Pfeiffer et al., 2024). Future research should furthermore investigate the possibilities of Fm theta NFT by incorporating implicit covert NFT protocols to eliminate confounding effects of traditional NFT learning paradigms (Muñoz-Moldes & Cleeremans, 2020). It should also be investigated what potential applications exist for using NFT to specifically induce a rebound effect of Fm theta activity and whether this effect could be intensified by using upregulation NFT.

## Conclusion

In conclusion, our findings provide new evidence for the notion that Fm theta downregulation NFT through traditional neurofeedback learning paradigms is more complex in comparison to other EEG frequency bands, yet its feasibility to induce a temporally limited subsequent reduction of task-related Fm theta power is reflected by our results. As our study did not show any significant NFT-related changes in task performance measures, we are not able to provide a definitive recommendation regarding the use of Fm theta downregulation for the improvement of volleyball setting anticipation at this point. Nevertheless, the ability to achieve a reduction of Fm theta activity during a complex cognitive task following theta NFT holds promise for future applications in the domain of sports and beyond.

## Acknowledgements

The authors would like to thank the Field of Excellence COLIBRI (Complexity of Life in Basic Research and Innovation, University of Graz) and Rebecca Teuschl for her assistance during data acquisition. This study was financially supported by the University of Graz for coverage of publication fees. No additional external funding was received for this study.

## Author Contributions

Conceptualization: Thomas Kanatschnig, Norbert Schrapf, Markus Tilp, Silvia Erika Kober. Data curation: Thomas Kanatschnig.

Formal analysis: Thomas Kanatschnig. Investigation: Thomas Kanatschnig.

Methodology: Thomas Kanatschnig, Lisa Maria Berger, Silvia Erika Kober. Project administration: Thomas Kanatschnig, Silvia Erika Kober.

Resources: Norbert Schrapf. Software: Lisa Maria Berger.

Supervision: Markus Tilp, Silvia Erika Kober. Validation: Thomas Kanatschnig.

Visualization: Thomas Kanatschnig.

Writing – original draft: Thomas Kanatschnig.

Writing – review & editing: Thomas Kanatschnig, Lisa Maria Berger, Norbert Schrapf, Markus Tilp, Silvia Erika Kober.

## Supplementary Material

**Fig. S1.**
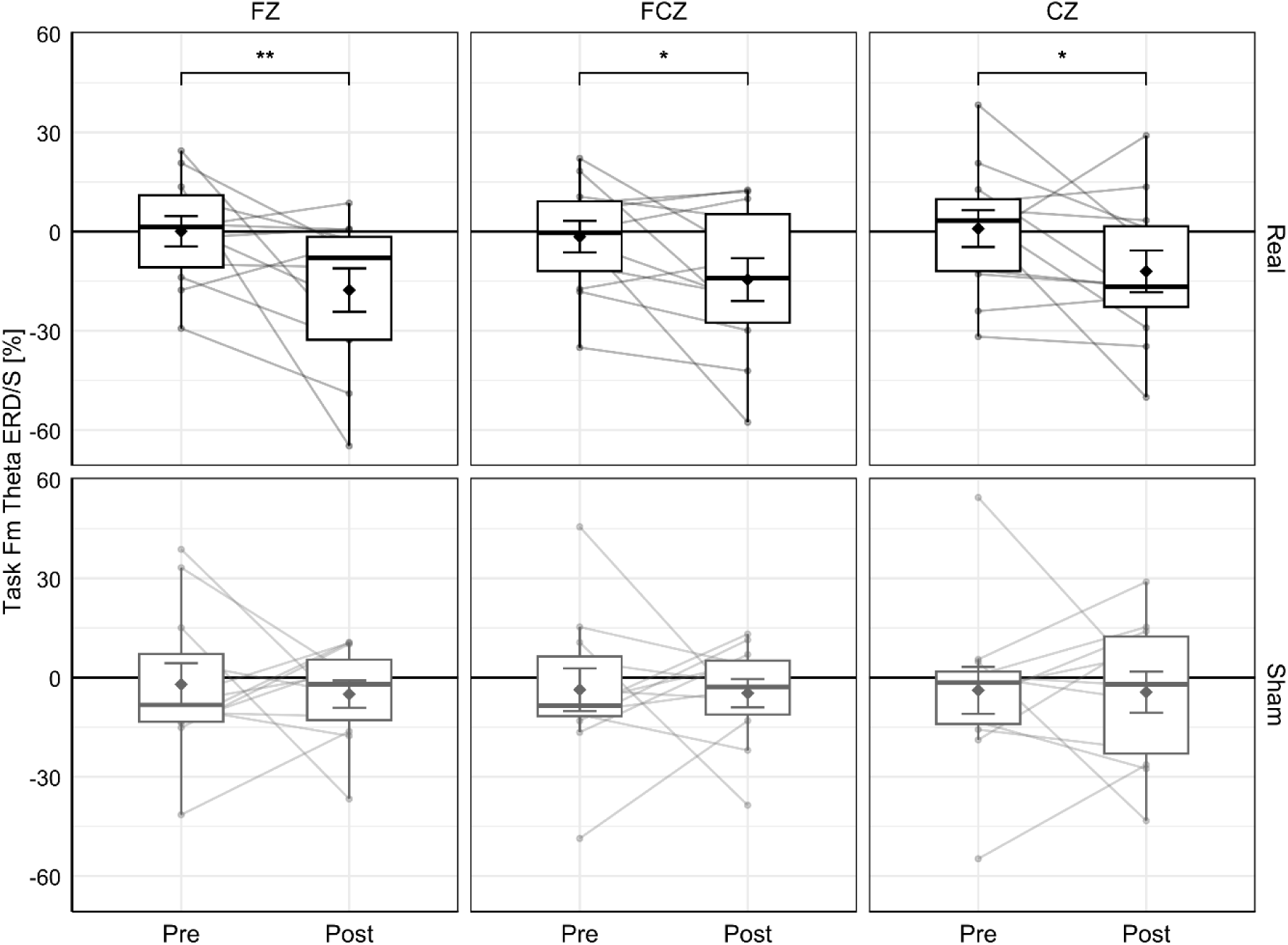
Results for the analysis of Task Fm Theta ERD/S separated by ROI electrodes. Pre/post differences of individual participants are represented by line-connected dots. Means and standard errors are represented within boxplots by diamond symbols and error bars. Significant pre/post differences, as determined through LMM analysis, are indicated with asterisks (*: *p* < .05; **: *p* < .01).

**Table S1.** Deviations from preregistration.

| Method: | Prereg: | Article: |
| --- | --- | --- |
| Dependent variable | ... For the analysis of fm-theta ERD/S during the task, a pre-stimulus “Baseline” phase (2 s visual fixation) will act as a reference to the “Video” phase (last 3 s before the end of each video stimulus), as well as the “Freeze” phase (0.5 s freeze of last video frame immediately before response), which are the main timeframes of interest. ... | In the present article the task has been renamed from “tactical decision-making task” into “volleyball setting anticipation task” (VSAT). As the definitions of specific phases of the task have been expanded over time, what was previously referred to as the “Video” phase was renamed to “Rally” phase in the present article, analogously to our previous article in which neurophysiological substrates of volleyball anticipation were investigated (Kanatschnig et al., 2025). As these deviations are merely semantical, they did not impact our statistical analyses. |
| Analyses | ... The analysis of behavioral performance and brain activity during the tactical decision-making task, will be conducted by means of four separate mixed-design ANOVAs. These will consist of Group and Time as independent variables and task accuracy and task response time, as well as fm-theta ERD/S, during the Video and Freeze phases, as dependent variables, including Holm-corrected post-hoc t-test comparisons where necessary. ... | With the use of LMM we deviated from our preregistered methods, in which we originally planned to use mixed-design ANOVA. However, the use of a more comprehensive LMM structure can be deemed an improvement of test severity, which makes this deviation an overall beneficial change regarding statistical rigor (Lakens, 2024). In the preregistration we also specified to investigate the Freeze phase of the task. Fm Theta ERD/S during the Freeze phase was analyzed analogously to the Rally phase. However, due to a lack of relevant additional information, we decided to omit the results of the Freeze phase in the main article, yet they are available for inspection through the provided online materials (https://osf.io/bu924/; Kanatschnig et al., 2024). |

*Note.* Methods that were preregistered for this study at the Open Science Framework (https://osf.io/hzg9j), for which the actual implementation in the article deviated are described under Prereg. Implemented procedures are described under Article.

**Table S2.** Comparative statistics for main sociodemographic and control variables.

| **Variables:** | **Real (*n* = 12):** | **Sham (*n* = 12):** | **Statistics:** |  |  |  |
| --- | --- | --- | --- | --- | --- | --- |
| **Categorical** | | | *df* | $\chi^2$ | *p* | *V* |
| Sex (female/male) | 8/4 | 8/4 | 1 | 0 | .999 | .00 |
| “Do you currently play actively in a volleyball club?” (no/yes) | 12/0 | 12/0 | 1 | 0 | .999 | .00 |
| “Do you play volleyball in your free time?” (no/yes) | 6/6 | 11/1 | 1 | 3.23 | .072 | .37 |
| “Are you currently attending one or more volleyball courses at the University Sports Institute (USI)?” (no/yes) | 12/0 | 12/0 | 1 | 0 | .999 | .00 |
| “Apart from volleyball, do you play sports regularly?” (no/yes) | 2/10 | 1/11 | 1 | 0 | .999 | .00 |
| **Continuous** |  |  | *df* | *t* | *p* | *d* |
| Age in years | 24.00 (±5.24) | 26.33 (±3.47) | 19.1 | -1.29 | .214 | -0.53 |
| “On average, how many hours a week do you play volleyball? (Please provide an approximate estimate of the number of hours.)” | 0.43 (±0.69) | 0.08 (±0.29) | 14.7 | 1.59 | .132 | 0.65 |
| “Approximately how many years of volleyball experience do you have? (Please provide a rough estimate of the years.)” | 2.33 (±3.06) | 1.83 (±2.82) | 21.9 | 0.42 | .681 | 0.17 |
| “On average, how many hours per week do you exercise? (Please provide an approximate estimate of the number of hours.)” | 3.33 (±2.05) | 5.67 (±4.42) | 15.5 | -1.66 | .117 | -0.68 |
| EHI | 74.58 (±17.38) | 77.08 (±21.89) | 20.9 | -0.31 | .760 | -0.13 |

*Note.* Absolute participant counts for respective answer options are presented for each categorical variable, while means and standard deviations are presented for continuous variables, within the respective group columns. Categorical variables were examined using $\chi^2$ -tests of independence plus Cramer's *V* calculations, while continuous variables were examined using independent samples *t*-tests plus Cohen's *d* calculations. Degrees of freedom for independent samples *t*-tests were adjusted by means of the Welch method (Welch, 1947); “EHI”: Edinburgh Handedness Inventory (Oldfield, 1971).

**Table S3.** Comparative statistics for visual analog scale variables.

| Variables: | Real (*n* = 12): | Sham (*n* = 12): | Statistics: |  |  |  |
| --- | --- | --- | --- | --- | --- | --- |
| Visual analog scale (VAS) |  |  | *df* | *t* | *p* | *d* |
| 1. “How difficult did you find it to predict the pass position before you completed the neurofeedback training?” (Not difficult at all/Very difficult) | 60.92 (±21.85) | 75.83 (±19.97) | 21.8 | -1.75 | .095 | -0.71 |
| 2. “How difficult did you find it to predict the pass position after you completed the neurofeedback training?” (Not difficult at all/Very difficult) | 52.67 (±19.54) | 65.00 (±26.57) | 20.2 | -1.30 | .210 | -0.53 |
| 3. “How difficult did you find it to raise the middle bar during neurofeedback training?” (Not difficult at all/Very difficult) | 50.83 (±31.97) | 68.75 (±24.64) | 20.7 | -1.54 | .139 | -0.63 |
| 4. “How difficult did you find it to keep the two outer bars as low as possible during the neurofeedback training?” (Not difficult at all/Very difficult) | 29.83 (±28.23) | 28.92 (±29.83) | 21.9 | 0.08 | .939 | 0.03 |
| 5. “How do you think your ability to regulate your own brain activity has changed over the course of the neurofeedback training?” (No improvement/Strong improvement) | 50.25 (±23.13) | 35.58 (±23.91) | 22.0 | 1.53 | .141 | 0.62 |
| 6. “How motivated were you to complete the prediction tasks and the neurofeedback training?” (Not motivated at all/Very motivated) | 81.92 (±14.35) | 81.75 (±13.42) | 21.9 | 0.03 | .977 | 0.01 |
| 7. “How has your motivation changed over the course of testing?” (Decreased significantly/Increased significantly) | 50.33 (±19.50) | 54.00 (±24.16) | 21.1 | -0.41 | .687 | -0.17 |
| 8. “How concentrated were you during the prediction tasks and the neurofeedback training?” (Not concentrated at all/Very concentrated) | 71.58 (±16.03) | 77.75 (±20.11) | 21.0 | -0.83 | .416 | -0.34 |
| 9. “How has your concentration changed over the course of testing?” (Decreased significantly/Increased significantly) | 50.42 (±24.70) | 52.33 (±26.09) | 21.9 | -0.19 | .855 | -0.08 |
| 10. “Were you able to recognize patterns in the game scenes during the prediction tasks that helped you with the tasks?” (Not at all/Very much) | 53.83 (±25.21) | 42.00 (±22.25) | 21.7 | 1.22 | .236 | 0.50 |
| 11. “How much do you think your previous knowledge of volleyball helped you with the prediction tasks?” (Not at all/Very much) | 17.42 (±14.18) | 21.58 (±21.77) | 18.9 | -0.56 | .585 | -0.23 |
| 12. “In the prediction tasks, did you feel more like guessing or knowing the answer?” (Guess/Know) | 39.83 (±23.98) | 34.33 (±25.21) | 21.9 | 0.55 | .590 | 0.22 |

*Note.* Means and standard deviations for VAS variables are presented within the respective group columns. Labels for the respective low (0) and high (100) ends of each scale are provided with each VAS variable. Categorical variables were examined using $\chi^2$ -tests of independence plus Cramer's *V* calculations, while continuous variables were examined using independent samples *t*-tests plus Cohen's *d* calculations. Degrees of freedom for independent samples *t*-tests were adjusted by means of the Welch method (Welch, 1947).

**Table S4.** Subject-level statistics and neurofeedback strategies.

| **Code:** | **Group:** | **NFT Fm Theta**<br>**ERD/S [slope]:** | **Task Fm Theta**<br>**ERD/S [diff %]:** | **Rest Fm Theta**<br>**[diff log($\mu V^2$)]:** | **NFT strategies:** |
| --- | --- | --- | --- | --- | --- |
| 1 | Real | 0.20 | -62.48 | 0.60 | nothing/relax, concentration bar, internal speech, concentration body, motor imagery |
| 2 | Real | 3.41 | -9.89 | 0.10 | concentration bar, nothing/relax, internal speech |
| 3 | Real | -5.53 | 13.19 | -0.09 | concentration internal, visual imagination |
| 5 | Real | 16.20 | -45.19 | 0.05 | concentration bar |
| 6 | Real | -4.22 | -8.79 | -0.28 | visual imagination |
| 7 | Sham | -0.29 | -9.78 | 0.05 | motor imagery, nothing/relax, concentration bar |
| 8 | Sham | 6.06 | 23.88 | 0.07 | nothing/relax, concentration bar |
| 9 | Sham | -0.36 | 20.69 | -0.12 | concentration bar, internal speech, moving eyes up/down |
| 10 | Sham | -1.33 | 22.99 | -0.25 | motor imagery, concentration internal, internal speech, body tension/relaxation |
| 11 | Sham | -0.25 | 3.24 | 0.28 | concentration internal, internal speech, emotion |
| 12 | Sham | -1.81 | -4.81 | -0.07 | concentration internal |
| 13 | Real | 0.22 | 6.37 | 0.18 | concentration bar, visual imagination |
| 14 | Real | -3.24 | -2.92 | -0.03 | concentration bar |
| 15 | Real | -1.55 | 4.55 | -0.13 | visual imagination, internal speech |
| 16 | Real | 3.54 | -24.66 | -0.40 | concentration bar |
| 17 | Real | 0.93 | -32.24 | -0.35 | nothing/relax |
| 18 | Real | -3.82 | -7.53 | -0.38 | motor imagery, nothing/relax |
| 19 | Sham | -1.85 | -53.79 | -0.09 | concentration bar, internal speech |
| 20 | Sham | -0.25 | -49.67 | -0.07 | concentration bar |
| 21 | Sham | 4.15 | -8.69 | 0.47 | nothing/relax, motor imagery, slow breathing |
| 22 | Sham | 3.20 | 29.74 | 0.60 | visual imagination |
| 23 | Sham | -2.31 | -7.96 | 0.21 | slow breathing, concentration bar, body tension/relaxation |
| 24 | Real | 3.52 | -5.21 | 0.15 | concentration internal, emotion |
| 25 | Sham | -2.63 | 15.89 | -0.07 | concentration bar, visual imagination |

*Note.* Neurophysiological statistics are presented for each participant (nr. 4 was excluded) alongside reported NFT strategies categorized by common content. Values for NFT Fm Theta ERD/S represent NFT slopes, while Task Fm Theta ERD/S and Rest Fm Theta ERD/S represent pre-/post-NFT differences.

## Footnotes

1 Certain panels in Fig. 1A have been removed from this preprint in accordance with visualization requirements. An unredacted version is intended for inclusion in the peer-reviewed publication.

